# *Wolbachia* strain *w*Au efficiently blocks arbovirus transmission in *Aedes albopictus*

**DOI:** 10.1101/844670

**Authors:** Maria Vittoria Mancini, Christie S. Herd, Thomas H. Ant, Shivan M. Murdochy, Steven P. Sinkins

## Abstract

The global incidence of arboviral diseases transmitted by *Aedes* mosquitoes, including dengue, chikungunya, yellow fever, and Zika, has increased dramatically in recent decades. The release of *Aedes aegypti* carrying the maternally inherited symbiont *Wolbachia* as an intervention to control arboviruses is being trialled in several countries. However, these efforts are compromised in many endemic regions due to the co-localization of the secondary vector *Aedes albopictus*, the Asian tiger mosquito. *Ae. albopictus* has an expanding global distribution following incursions into a number of new territories. To date, only the *w*Mel and *w*Pip strains of *Wolbachia* have been reported to be transferred into and characterized in this vector. A *Wolbachia* strain naturally infecting *Drosophila simulans*, *w*Au, was selected for transfer into a Malaysian *Ae. albopictus* line to create a novel triple-strain infection. The newly generated line showed self-compatibility, moderate fitness cost and complete resistance to Zika and dengue infections.

**Author summary:** *Aedes albopictus*, the invasive Asian tiger mosquito, is responsible for numerous outbreaks of important viruses such as dengue and Zika in various regions of the world. The need for alterative control interventions propelled the development of a novel approach that exploits a natural insect symbiont, *Wolbachia*; when transferred into non-native hosts, these maternally-inherited bacteria are able to interfere with the transmission of mosquito-borne viruses, and also provide reproductive advantages to the host, offering a promising tool for self-sustaining field applications. Currently, several field trials are ongoing for the primary vector of dengue and several other arboviruses, *Aedes aegypti*, providing encouraging results. In this study, *Ae. albopictus* has been targeted for a similar approach: this mosquito species naturally carries two strains of *Wolbachia*. The artificial introduction of a third, non-native strain made this line less able to transmit dengue and Zika viruses and had an impact on its fitness.

## Introduction

The Asian tiger mosquito *Aedes albopictus* is an increasingly prominent vector of arboviruses. This anthropophilic and peridomestic mosquito species is an aggressive daytime biter with a capacity to invade both temperate and tropical areas by adapting to different climates and producing overwintering eggs. It originates in Asia, but is now widely distributed across Europe, Africa, the Americas and the Pacific, and the identified transboundary mechanisms involved in its expansion contribute to a high invasiveness and ever-increasing geographic range (1, 2).

*Ae. albopictus* has been incriminated as a vector of more than 20 arboviruses (3); it acted as the primary vector of some lineages of chikungunya virus (CHIKV) during outbreaks in La Reunion (4, 5) and Italy (6). It is also a well-characterized secondary vector of dengue virus (DENV) (7), with similar competence to the primary vector *Ae. aegypti* (although a lower propensity to bite humans), and is responsible for autochthonous DENV transmission in Europe (8, 9). Furthermore, in addition to a capacity to be artificially infected with Zika virus (ZIKV) in a laboratory setting (10), field-collected *Ae. albopictus* in the Americas were recently found to be Zika-positive (11).

Given the enormous impact on public health, and economic impacts imposed by the management of outbreaks in terms of direct and indirect medical costs, there is a need for investment in arbovirus prevention programmes (12, 13). Due to the compromised efficacy and sustainability of traditional vector control approaches – mainly based on the use of insecticides-novel control strategies are being explored, such as biological control using transmission-blocking symbionts. Among the most promising candidates are *Wolbachia*, maternally inherited intracellular alpha-proteobacteria commonly found in a wide variety of arthropods; several strains have been shown to provide resistance to different RNA-viruses (14–18), especially when artificially transferred into non-native hosts (19, 20). Many strains of *Wolbachia* induce a reproductive distortion in the host known as cytoplasmic incompatibility (CI)(21), which occurs when the sperm of *Wolbachia*-carrying males is modified, resulting in early embryonic death when these sperm fertilize eggs from non-carrier females. In contrast, females carrying the same *Wolbachia* strain are able to ‘rescue’ the sperm modification, producing viable offspring.

*Ae. albopictus* naturally carries two strains of *Wolbachia*, *w*AlbA and *w*AlbB (22), although at a relatively low overall density and mostly confined to the reproductive organs. There is evidence that density and bacterial tissue tropism play a role in *Wolbachia* virus-blocking ability. In fact, when *w*AlbB is transferred from *Ae. albopictus* into the non-native host *Ae. aegypti*, its density and tissue tropism increase, together with the ability to protect the host against arboviruses (17, 23).

Several strains of *Wolbachia* have been characterized in different hosts, revealing considerable diversity in terms of intracellular density, fitness effects, susceptibility to high temperatures, expression of CI and pathogen inhibition (24–27). The attributes of each strain within a novel host must be thoroughly characterised to ensure optimal deployment *Wolbachia* for disease control; there may be a balance between the level of virus blocking and host fitness costs, which will increase the threshold frequency that must be exceeded before it will spread and become established in populations. A stable *Ae. albopictus* line carrying the *w*Mel strain, transferred from its natural host *Drosophila melanogaster*, showed reduced vector competence for DENV and CHIKV(28, 29). However, this strain has been shown to be susceptible to density reductions and maternal leakage in *Ae. aegypti*, when larval stages were exposed to high temperatures (24, 30).

A *Wolbachia* strain from *Drosophila simulans*, *w*Au, has been stably transferred into *Ae. aegypti*, to which it provides unusually high level of inhibition of Semliki Forest Virus (SFV), DENV and ZIKV; additionally, there is evidence that this strain is less susceptible to heat-stress more than *w*Mel (17). Similarly in its natural host, this strain reaches high densities and protects against a wide range of *Drosophila* viruses. Although it does not induce CI, spread to relatively high frequency has been reported in field populations of *D. simulans* in Australia; this may be a consequence of increased host fitness (31). In light of this, *w*Au was selected for transfer into a Malaysian *Ae. albopictus* line to create a novel triple-strain infection and characterized for parameters relevant to its potential field application for disease control.

## Methods

### Mosquito rearing

The *Ae*. *albopictus* wild-type strain (JF), origin Jalan Fletcher area of Kuala Lumpur, Malaysia, and maintained for >20 generations, was kindly provided by W.A. Nazni, Institute for Medical Research, Kuala Lumpur. A *w*Au-infected *Ae. aegypti* line (17) was used as the *Wolbachia* source. The *w*Mel single-infected *Ae. albopictus* line was generated as previously described (28). Mosquito colonies were maintained at standard 27°C and 70% relative humidity with a 12-hour light/dark cycle. Larvae were fed on tropical fish pellets (Tetramin, Tetra, Melle, Germany) and adults were offered 5% sucrose solution *ad libitum*. Blood meals were provided using a Hemotek artificial blood-feeding system (Hemotek, UK) using defribrinated sheep blood (TCS Biosciences, UK). Eggs were collected on wet filter paper (Grade 1 filter paper, Whatman plc, GE healthcare, UK) and desiccated for 5–10 days before hatching in de-ionized water containing 1g/L bovine liver powder (MP Biomedicals, Santa Ana, California, USA).

### Generation of JF.Neg and JF.*w*Au (*w*AlbA-*w*AlbB-*w*Au) lines

The *Wolbachia*-free JF.Neg was generated by treating adults with 1.25 mg/ml tetracycline dissolved in 5% sugar solution (32). After 4 generations, antibiotic-treated mosquitoes were 100% negative when analysed by qPCR, with no impact on general fitness, egg hatchability, larvae mortality and fecundity.

The generation of a triple-strain (*w*AlbA*-w*AlbB*-w*Au) line in *Ae. albopictus* involved the transfer of cytoplasm from embryos of the *Ae. aegypti* line carrying *w*Au; microinjections were performed as previously described (17). Female survivors (G_0_) were back-crossed to JF.Neg males, blood-fed and separated individually for oviposition. Strain specific-PCRs for target *Wolbachia* strains were performed, and only eggs from positive G_0_ were hatched. Progeny were also assayed for *Wolbachia* using PCR to assess germ-line transmission. Positive females were backcrossed with JF.Neg males and their progeny were selected until G_5_, when the line was self-compatible and stable for maintenance.

### *Wolbachia* strain-specific PCR and qPCR in whole mosquitoes and tissues

gDNA was extracted from 5-day old whole mosquitoes and tissues (ovaries, guts and salivary glands) using STE buffer (10uM Tris HCL pH 8, 100mm NaCl, 1mm EDTA) and diluted to 100 ng/μl using a NanoDrop spectrophotometer (Thermo Scientific, Waltham, Massachusetts, USA). Strain specific PCR was used for the screening of the newly generated line. Primers sequences are summarized in Table S1.

PCR reactions were set up using 1x Taqmaster mix (Vazyme) according to the manufacturer’s protocol. DNA was amplified with an initial denaturation at 94 °C for 3 min, followed by 30 cycles consisting of denaturation at 94 °C for 30 s, annealing at 55 °C for 30 s, extension at 72 °C for 30s, and a final step at 72 °C for 10 min. Samples were then visualized on a 1% agarose gel stained with SYBR safe DNA Gel Stain (ThermoScientific, UK).

*Wolbachia* density was assessed by qPCR using the relative quantification of the *Wolbachia* surface protein (*wsp*) gene against the homothorax gene (HTH) as reference gene. To specifically quantify the *w*AlbA, *w*AlbB, and *w*Au strains, and for the measurement of total *Wolbachia* density the following primers were used: *w*AlbA – qAlbAF and qAlbAR; *w*AlbB - 183F and QBrev2; *w*Au– *w*AuT3F and wAuT3R; wsp-WSP-F and WSP-R. The following program was used to run the qPCRs: 95 °C for 5 min, 40× cycles of 95 °C for 15 s and 60 °C for 30 s, followed by a melt-curve analysis. A Rotor Gene Q (Qiagen) was used with 2x QuantiNova SYBR.

### Adult Longevity

A cohort of 25 females and 25 males of JF.*w*Au, JF.Neg and JF was used for assessing adult survival. Individuals were maintained in 24.5×24.5×24.5cm rearing cages in the insectary, under standardized control conditions. Cages were blood-fed once a week and eggs were collected regularly. Mortality was monitored daily until no individuals were alive. Three biological replicates were performed.

### Fecundity and egg survival

Fecundity rate and egg survival were assessed using blood fed, fully engorged JF.*w*Au females. JF and *w*Au-*Ae.aegypti* were included as controls. 20 individuals were individualized inside up-turned cups on top of filter paper. After 3 days, the number of eggs laid per female was counted under a stereoscope.

Additionally, the impact of desiccation on egg survival was measured on collected eggs from blood-fed 5-day old females. Sections of filter papers with 300-400 eggs were cut and stored at 27°C and 70% relative humidity. After 5, 15, 30, 50 days post oviposition, sections were floated in water containing 1g/L bovine liver powder. Hatch rates were assessed by counting hatched eggs and L3-L4 instar larvae.

### Maternal inheritance and CI

CI induction between crosses with JF, JF.Neg and JF.*w*Au was evaluated between 25 males and 25 females in separated cages. Females were offered a blood meal and individualised for oviposition. Eggs were collected, desiccated for 5 days, counted and then, hatched in water containing 1 g/l bovine liver powder. Hatched larvae were counted to estimate egg hatch rates.

Maternal transmission of each of the three *Wolbachia* strains present (*w*AlbA, *w*AlbB and *w*Au) was evaluated by backcrossing JF.wAu females with JF.Neg males. Females were blood-fed and individualized for oviposition. Eggs, collected on damp filter papers were desiccated for 5 days and added to water for hatching. A selection of progeny was randomly sampled and analysed with strain-specific PCR.

### Fluorescent *in situ-*hybridization

Ovaries were dissected from 5 day old females using sterile forceps and needles in a drop of sterile PBS buffer for *Fluorescent In Situ Hybridization* (FISH) and then transferred in Carnoy’s buffer (chloroform:ethanol:acetic acid, 6:3:1) and fixed at 4°C overnight. Samples were then rinsed in PBS and incubated in a hybridization buffer containing 50% formamide, 25% 20xSSC, 0.2% (w/v) Dextran Sulphate, 2.5% Herring Sperm DNA, 1% (w/v) tRNA, 0.015% (w/v) DTT, 1% Denhardt’s solution, and 100 ng/ml of each probe. General *Wolbachia* probes annealing on the *wsp* gene were used as described before (16).

Samples were left to hybridize overnight in a dark-humid box at 37°C, washed twice in a solution containing 5% 20xSSC, 0.015% (w/v) DTT, and twice in a solution of 2.5% SSC, 0.015% (w/v) DTT in dH2O, at 55°C for 20 minutes. Tissues were then placed on a slide containing a drop of VECTASHIELD Antifade Mounting Medium with DAPI (Vector Laboratories, California, USA) and were visualized using a Zeiss LSM 880 confocal microscope (Zeiss, Oberkochen, Germany).

### Heat-stress response

Eggs from wild-type colonies, JF.*w*Au and *w*Mel-carrying *Ae. albopictus* were hatched and separated into experimental groups: larval density (200 larvae per 500 mL of water) and food were consistent between conditions. Heat-challenged larvae were maintained in Panasonic MLR-352-H Plant Growth Chamber incubator (Panasonic, Osaka, Japan). The temperature regime replicated in cabinets was a diurnal cycle of 27-37°C, while control larvae were reared at constant 27°C. Adults were all maintained at constant 27°C. *Wolbachia* density was quantified by qPCR on whole-bodies of 5-day old females and males, using *wsp* primers and *w*Au-specific primers.

### Virus challenge

7-day old JF.*w*Au and JF females were blood-fed with human blood, 5mM of phagostimulant ATP and virus suspension. Dengue virus was serotype 2 (New Guinea C Strain) at the final concentration in the blood meal of 10^8^ FFU/ml, while Zika virus was strain MP1751, obtained from Public Health England culture collections, at a final concentration in the blood-meal of 4×10^7^ FFU/ml. Engorged females were transferred in a climactic chamber at 27 C and 70% humidity for 14 days with access to 5% sucrose-soaked cotton wool. At the same time, a different cohort of females was anaesthetised on ice and injected with 10^9^ FFU/ml of DENV-2 in the thorax using a Nanoject II (Drummond Scientific, USA) hand-held microinjector. Similarly, 1.4×10^7^ FFU/ml of ZIKV virus was injected in the thorax of females.

After 14 days, whole-bodies of virus-injected females (DENV and ZIKV-infected) were homogenised with glass beads in Trizol (Sigma-Aldrich, MA, USA) on a Precellys 24 homogeniser (VWR).

Salivary glands of ZIKV blood-fed and heads of DENV blood-fed females were dissected in sterile conditions and transferred in Dulbecco’s Modified Eagle Medium (DMEM) medium supplemented with 2% fetal bovine serum (FBS), After being homogenized, the solution was transferred onto a monolayer of Vero cells for fluorescent focus assay (FFA). Primary antibody for Zika was a Mouse Monoclonal antibody DIII1B kindly provided by Prof. Arvind Patel (MRC-University of Glasgow-CVR); secondary antibody was the Goat anti-mouse Alexa Fluor 488, A-11001 (Thermo Scientific, Waltham, Massachusetts, USA). Celigo Imaging Cytometer (Nexcelom Bioscience, Lawrence, Massachusetts) was used for imaging plates.

Carcasses of dissected females were sampled in Trizol for quantification of ZIKV infection. RNA was extracted with Trizol according to manufacturer’s guidelines and diluted to 100ng/ul with ddH20; cDNA was synthesised All-In-One cDNA Synthesis SuperMix (Biotools, TX, USA). DENV-2 was quantified by qPCR using NS5-F and NS5-R set, while ZIKV 835 and ZIKV 911c primers were used for ZIKV quantification. Values were normalised to the RpS17 as reference by relative expression (Pfaffl method).

### Statistical analysis

Graphics were generated using the ‘ggplot2’ package of R Studio (RStudio Inc., Boston, Massachusetts, USA) of the R software (version 3.6.1). All statistical analyses were run using Prism version 7. Shapiro-Wilk Test was used for assessing normality distribution of data, and parametric and non-parametric tests were selected accordingly. Multiple comparisons were performed using Multiple t-tests and one-way ANOVA using Holm-Sivak’s or Bartlett’s tests. Survivals were statistically analysed using proportional hazard ratio model (Log-rank - Mantel-Cox test). Analysis of virus-challenged mosquitoes was performed using a Fisher’s exact test comparing rates of virus-positive and virus-negative samples.

## Results

### Strain generation and characterization

#### Maternal transmission and CI

The JF.*w*Au line was created by transferring cytoplasm from *w*Au-carrying *Ae. aegypti* into wild-type (*w*AlbA and *w*AlbB-carrying) *Ae. albopictus*, obtaining a stable triple-strain infection. From G_1_ to G_5_, females were backcrossed with JF males, blood-fed and individualized; the progeny were pooled only if the mother was positive for all the three strains. From G_5_, colonies were maintained without backcrossing and selecting females: random individuals tested positive in the following generations, indicating that the maternal vertical transmission of the three strains was complete under standard rearing conditions.

Crosses were set up in order to investigate patterns of compatibility and/or CI between the groups. As expected, *w*Au was not able to induce unidirectional CI when males of JF.*w*Au were crossed to wild-type females (JF). The JF.*w*Au line was self-compatible with a similar hatching rate when compared to wild-type. Complete CI was observed between *Wolbachia*-cured females mated with wild-type or triple-strain males, confirming that the *w*AlbA / *w*AlbB strains are still able to induce sterility in the presence of *w*Au.

**Table 1:** Crosses and CI patterns. *Wolbachia* triple-strain (JF.*w*Au), wild-type (JF) and tetracycline-treated *Ae. albopictus* (JF.Neg) were crossed to evaluate strains compatibility and CI phenotypes. Numbers represent percentages of hatch rates, while the total number of analysed eggs is shown in parentheses.

|  |  | Males |  |  |
| --- | --- | --- | --- | --- |
|  |  | JF | JF. wAu | JF.Neg |
| Females | JF | 79 (1106) | 57.3 (2093) | 51.6 (1176) |
|  | JF. wAu | 70 (2652) | 65.6 (1564) | 75 (1616) |
|  | JF.Neg | 0 (2039) | 0 (1838) | 81 (2051) |

As observed during line selection, JF.*w*Au showed complete maternal transmission of all three *Wolbachia* strains to progeny: *w*AlbA, *w*AlbB and *w*Au were found in 100% of the 100 individuals analysed among the progeny of individualized females.

#### *Wolbachia* density and tissue distribution

*Wolbachia* density from the whole-bodies of mosquitoes was quantified 5, 10 and 20 days post-eclosion (Fig.1A). Total *Wolbachia* in JF.*w*Au was found to be significantly higher than wild-type after 10 and 20 days post eclosion, similar to what was previously described in *w*Mel-single and triple infection in *Ae. albopictus* (28, 33). Nevertheless, when compared to *w*Au in *Ae. aegypti*, density is significantly lower, lending further support to previous findings that, when transferred into non-native hosts such as *Ae. aegypti*, *Wolbachia* undergo density modulations (17, 34). The density of the native *w*AlbA and *w*AlbB infections appeared to be unaffected by *w*Au co-infection. (Fig.1B).

There is evidence that the strength of viral blockage and the ability to be maternally inherited can also depend on *Wolbachia* tissue tropism (35). Total *Wolbachia* density was quantified on dissected ovaries, salivary glands and midguts (Fig. 1C): generally, the JF.*w*Au line showed relatively higher densities in somatic tissues, reaching very high density in the ovaries when compared to wild-type.

**Fig. 1.**
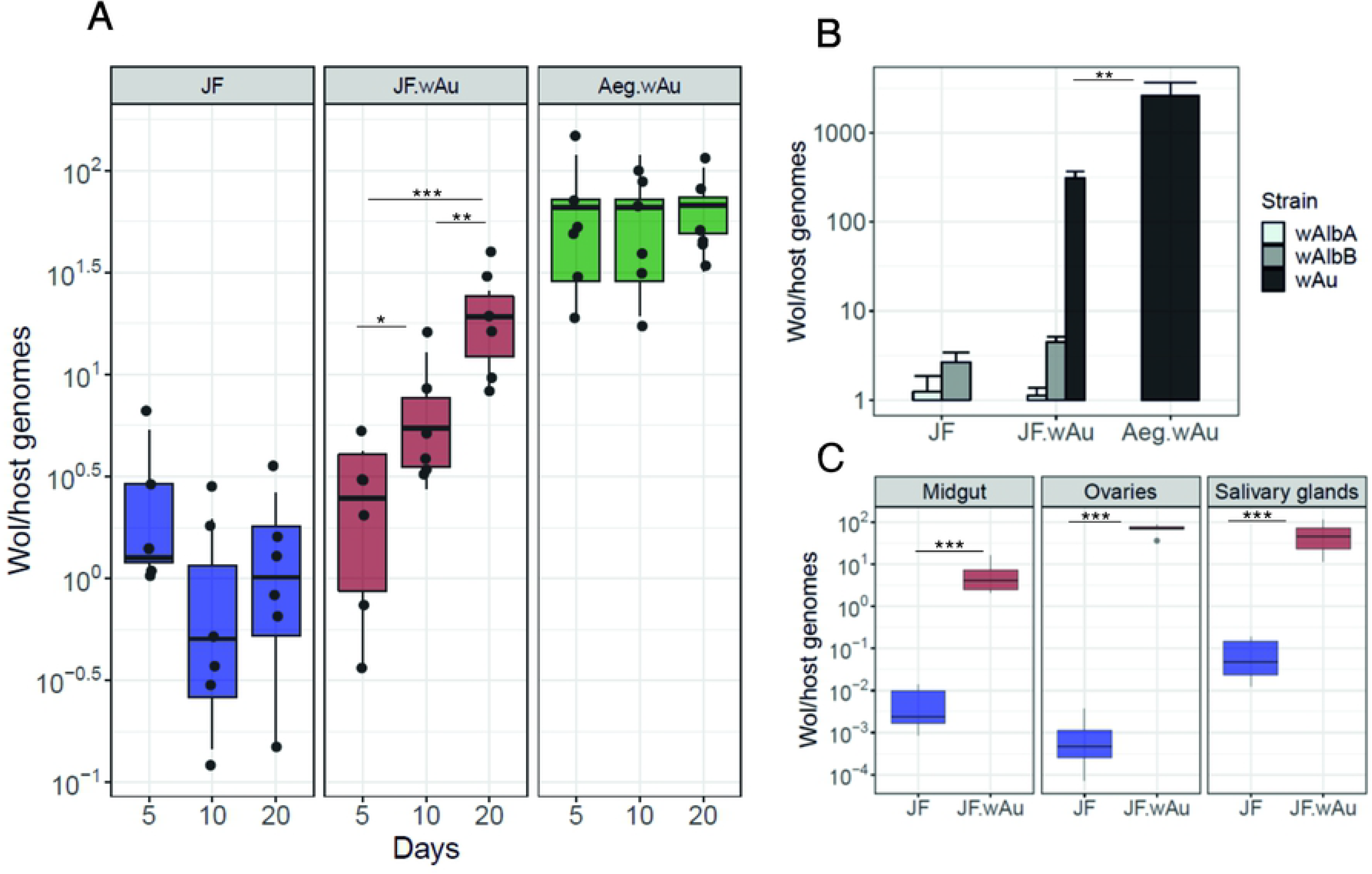
*Wolbachia* density and tissue distribution. A) 5, 10 and 15 day-old adults were tested for total *Wolbachia* density. Boxplots represent 6 biological replicates: the centre indicates the median of densities and whiskers represent upper and lower extremes. B) Strain-specific qPCR was used for assessing the density of each strain of the triple infection. Bars represent the average of 6 biological replicates. Error bars show SE. C) *Wolbachia* tissue distribution was measured in 6 pools of 3 sets of dissected ovaries, salivary glands and midguts. t-test and Mann-Whitney test were used for statistical analyses. *p<0.05,**p<0.001, ***p<00001. Non-significant differences are not indicated.

#### Fitness characterization

Additionally, several traits of the fitness of the newly generated line were characterized: adult lifespan, fecundity and fertility of females (Fig.2). Although significantly reduced when compared to wild-type, JF.*w*Au females showed a similar life span to the *Wolbachia*-cured *Ae. albopictus* (Fig.2A). The survival of JF.*w*Au males, on the contrary, was not affected by *w*Au and was not different from the wild-type strains (Fig. 2B). Fertility of individualized females was compared between JF.*w*Au, JF and *Ae. aegypti*-*w*Au showing no significant difference on the amount of laid eggs (Fig. 2C). In contrast, the co-infection of *w*Au seemed to have an impact on the hatching rate of desiccated eggs of the JF.*w*Au line. After 15 days, the number of hatched eggs notably dropped with respect to wild-type eggs, similarly to *Ae. aegypti*-*w*Au, further confirming that high density strains have an impact on egg quiescence (Fig. 2D).

**Fig 2.**
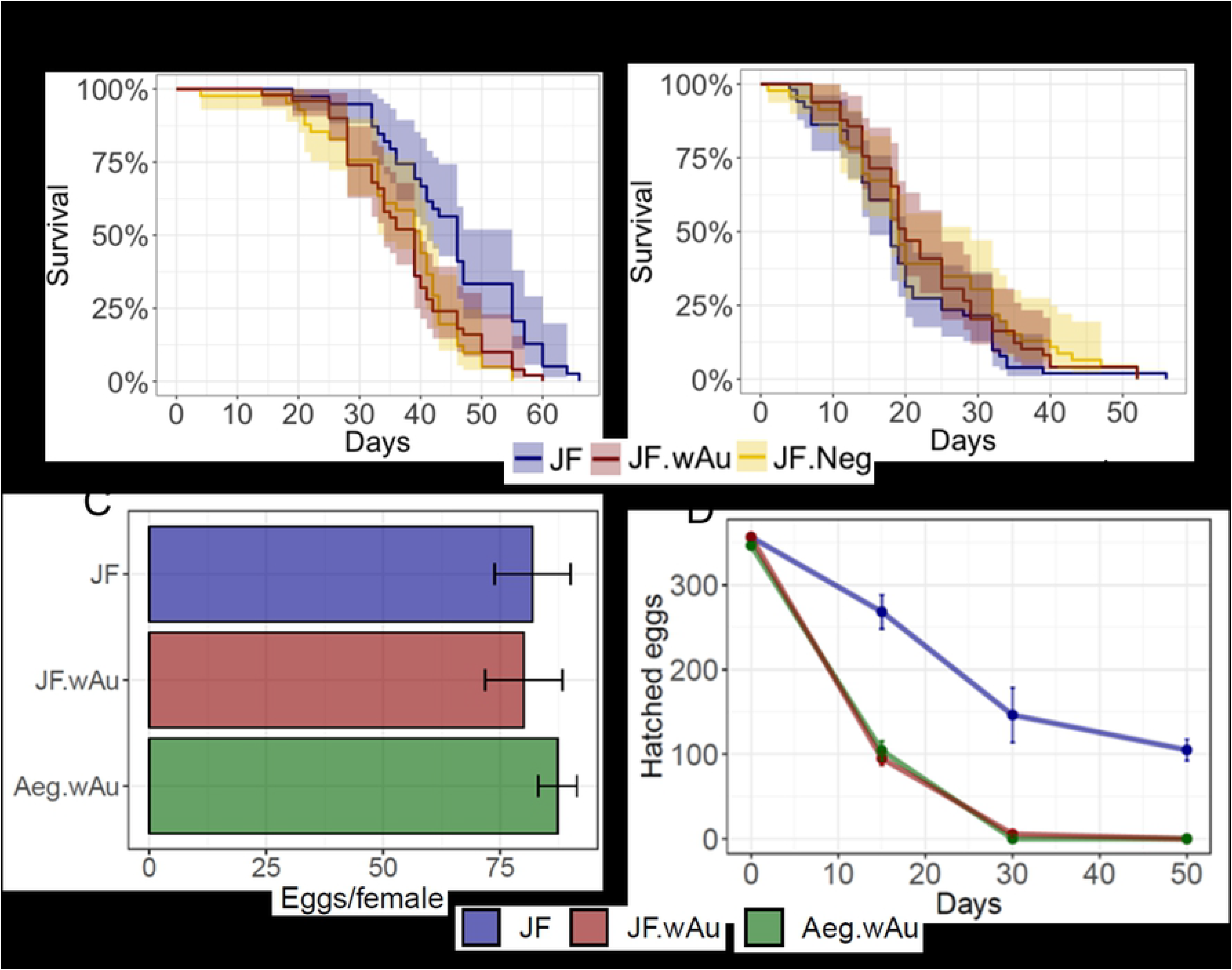
Fitness traits characterization. A-B) Females and males life span of JF, JF.wAu and JF.Neg was measured and compared using proportional hazard ratio model (Log-rank - Mantel-Cox test). C) Fecundity of lines determined by counting the number of eggs from individual females. SE is indicated. One-way ANOVA with Bartlett’s test was used for measuring statistical differences. D) Mean hatching rate over time. Statistical differences between datasets was measured using Two-way ANOVA with Dunnett’s comparison.

#### Effects of high temperature exposure

Exposure to high temperatures during larval development is known to reduce the density of some strains of *Wolbachia* in *Ae. aegypti*, including the *w*Mel strain (17, 24). Consequently, the effects of high temperature exposure were investigated in a previously generated *Ae. albopictus* line carrying only the *w*Mel strain (28, 29), and in the newly generated JF.*w*Au. Larvae were exposed to a diurnal cycle with fluctuations from 27 to 37°C in a programmable incubator. The general *Wolbachia* density in different lines was assessed by qPCR in 5-day old females and males. As observed in *Ae. aegypti*, the *w*Mel strain was particularly sensitive to exposure to high temperatures, showing a decrease in density of several orders of magnitude (Fig. 3).

**Fig. 3.**
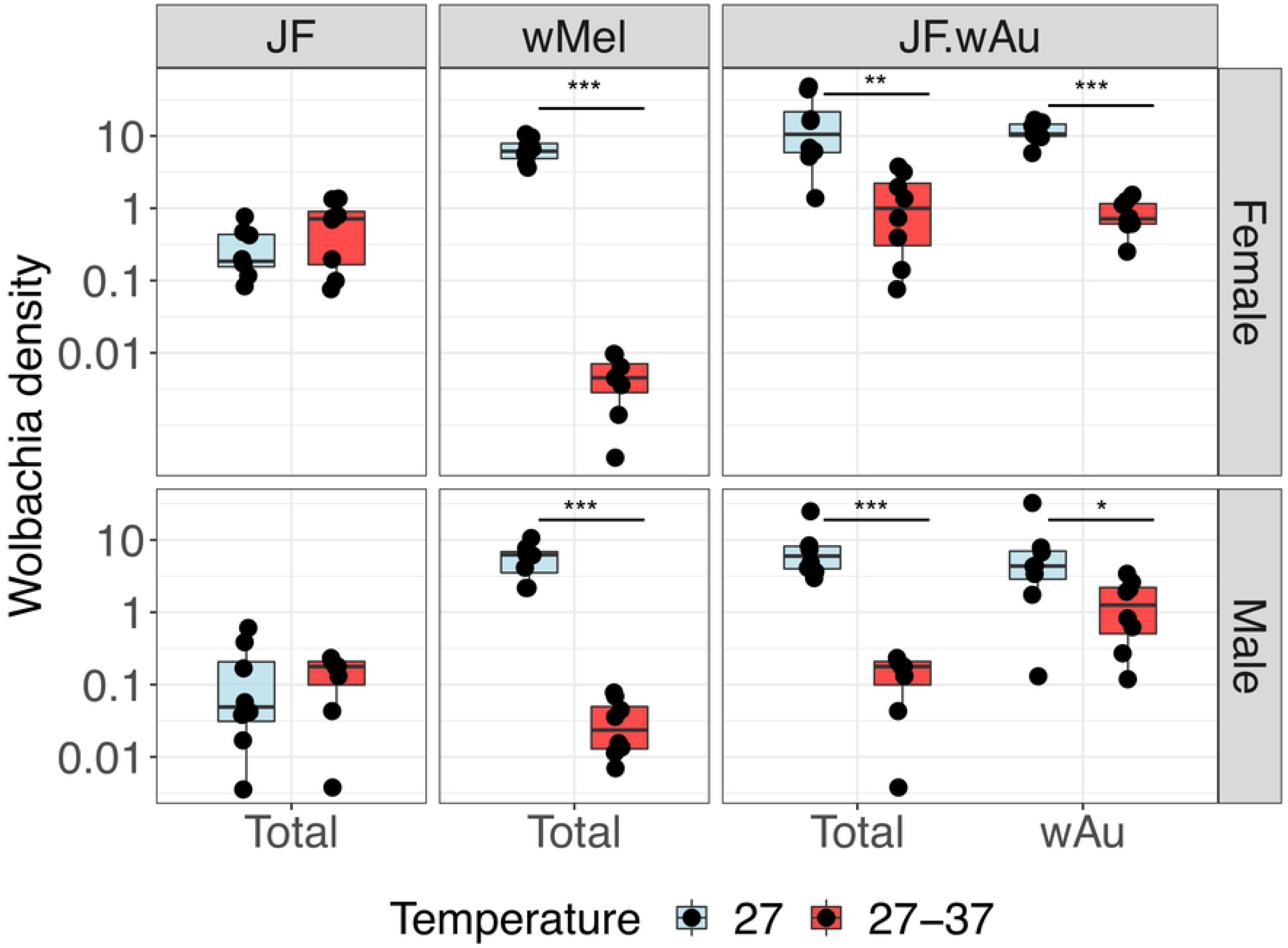
Exposure to high larval breeding temperatures. Larvae from *w*Mel-*Ae. albopictus*, JF.*w*Au and JF lines were reared with a diurnal cycle of 27-37°C. Controls were maintained at constant 27°C. Boxplots represent 8 biological replicates: the centre indicates the median of densities and whiskers represent upper and lower extremes. T-test and Mann-Whitney test were used for statistical analyses. *p<0.05,**p<0.001, ***p<00001. Non-significant differences are not indicated.

When transferred into *Ae. aegypti*, *w*Au was somewhat sensitive to thermal stress, and its density in adults underwent a significant decrease (17). Similarly, the total *Wolbachia* density of the JF.wAu lines displayed a drop in heat-treated individuals compared to controls maintained at constant 27°C, although mean density remained considerably higher than observed for *w*Mel (Fig. 3). The density of the native strains in *Ae. albopictus* wild-type, *w*AlbA and *w*AlbB, did not appear to be perturbed by exposure to high temperatures. Consequently, the specific contribution of *w*Au to the drop in density was further investigated: the quantitative analysis confirmed a significant decrease in *w*Au in females and males, suggesting a similar response to that observed in *Ae. aegypti*.

#### Virus inhibition

DENV-2 and ZIKV were used to evaluate the ability of this line to transmit arboviruses. Mosquitoes were infected through intra-thoracic injection of the virus or through an infected blood-meal. Whole bodies of blood-fed JF.*w*Au showed significantly reduced DENV-2 loads in whole carcasses (abdomens, heads and thoraxes), compared to JF, confirming the ability of the *w*Au strain to be an efficient viral blocker in *Ae. albopictus* (Fig.4A). This was further demonstrated when high viral loads (10^9^ FFU/ml) were intra-thoracically injected directly into the mosquito haemolymph, bypassing the natural midgut barrier: a significant reduction of DENV-2 genome copies was observed between the triple-strain line compared to wild-type (Fig.4A). A second independent DENV challenge (Fig. 4B) was performed: in this instance, after 14 days, mosquito heads and thoraxes were analysed by FFA in order to assess virus dissemination, while abdomens were analysed separately by qPCR. This replicate provided further validation, confirming the complete blockage of the virus in both tissues.

**Fig. 4.**
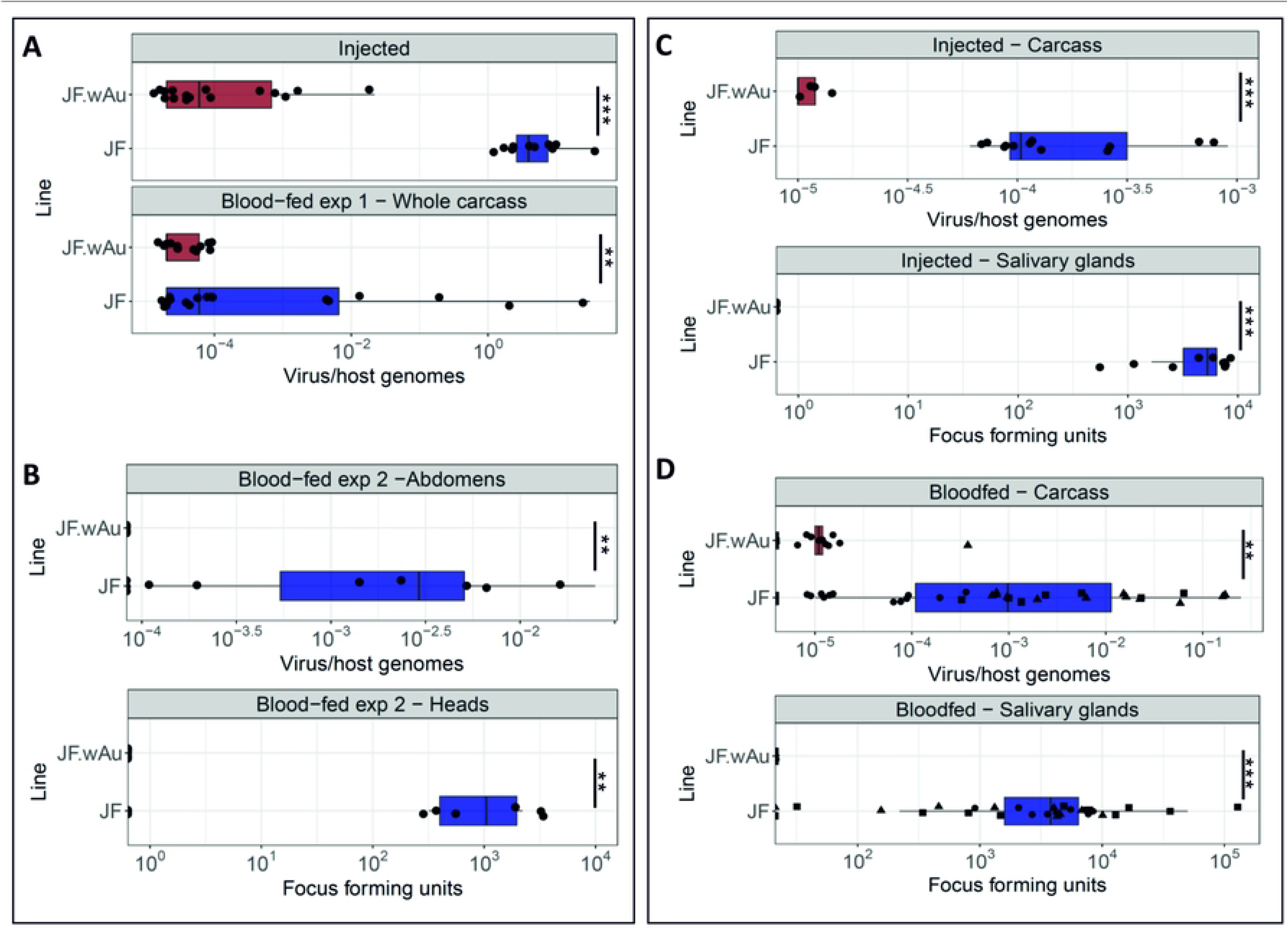
DENV2 and ZIKV challenges of JF.*w*Au and wild-type mosquitoes. A) DENV2 genome copies were quantified in whole bodies of 3-5 day old mosquitoes using qPCR on injected and blood-fed females after 14 days. B) Viral genome copies and foci number were quantified after a DENV2-infected blood meal on abdomens and heads. C) ZIKV genome copies (in carcasses) and viral foci number (in dissected salivary glands) of injected and D) blood-fed 3-5 day old mosquitoes. The latter graph summarises three independent biological replicates; different shapes represent data points from each replicate. p<0.001, ***p<00001.

The high blocking phenotype was found to be consistent when mosquitoes were challenged with ZIKV. Similar to the DENV challenges, even when very high viral titers were directly injected into the female hemolymph, the *w*Au strain is able to block the dissemination and the transmission of the virus (Fig 4C). Similarly, no virus was detected by qPCR on carcasses or by FFA on salivary glands of the JF.wAu line after three independent ZIKV-infected blood meals, compared to the wild-type (Fig 4D).

## Discussion

*Wolbachia-*based field interventions to control arboviruses are under investigation in a number of countries for *Ae. aegypti*, demonstrating promising efficacy for mosquito population suppression and replacement strategies. Nevertheless, dengue control in many endemic regions still remains a challenge due to the sympatric secondary vector *Ae. albopictus*. Its abundance in many urban and suburban spaces, and its low active dispersal activity makes this vector a suitable candidate for releases in confined urban sites of endemic regions (36, 37). Recently, the use of an artificial triple-*Wolbachia* infection combined with the sterile insect technique (SIT) for population suppression interventions demonstrated a successful reduction of *Ae. albopictus* population density in open field trials in China (38). Additionally, positive results of IIT (Incompatible Insect Technique) for population suppression were also obtained after preliminary trials in Italy of released of *Ae. albopictus* carrying the strain *w*Pip (39, 40).

As a promising option for population replacement transmission-blocking approaches, a line carrying only the *Ae. albopictus w*Mel strain was previously shown to produce no detectable fitness costs in this host, to induce bidirectional CI and to be refractory to DENV and CHICK in laboratory tests (28, 29). To develop a line with greater invasiveness, a triple-strain infection combining *w*Mel with the native strains, *w*AlbA and *w*AlbB, was also subsequently created with the aim of generating unidirectional CI: surprisingly however, the line was found to be self-incompatible and thus, not a practical option to be deployed in the field (33). Additionally, as previously demonstrated in *Ae. aegypti*, and now also here in *Ae. albopictus*, the exposure of *w*Mel-carrying larvae to high temperatures during rearing produces large decreases in overall *Wolbachia* density. The deployment of *w*Mel in *Ae. albopictus* for dengue control is therefore likely to be impractical in tropical areas where larval stages experience very high larval site temperatures. The exposure of JF.*w*Au to cyclical high temperatures during the larval stages resulted in a moderate impact on the overall density of this *Wolbachia* strain, in this respect offering a more suitable dengue control tool for use in hotter equatorial areas when compared to *w*Mel.

The novel JF.*w*Au line generated in this study was completely refractory to DENV and ZIKV, and *w*Au consistently demonstrated complete viral inhibition in both injected and blood-fed mosquitoes. JF.*w*Au displayed a higher overall *Wolbachia* density when compared to wild type, confirmed in the dissected midgut epithelia, salivary glands and ovaries; this higher overall density is mostly a result of the *w*Au strain, as was previously observed in *Ae. aegypti* (17), while the densities of the two native strains are comparable to their wild-type counterparts. The presence of *w*Au does not cause exclusion or significant density reduction of either of the native strains, ensuring complete maternal transmission of the three strains. In contrast to what was previously observed for the *w*Mel*w*AlbA*w*AlbB triple-infection, the JF.*w*Au line displayed full self-compatibility, and JF.*w*Au females were fully compatible when crossed with wild-type males.

The intricate pattern of interactions between different *Wolbachia* strains and their hosts determines the trade-offs between pathogen blocking, host fitness and invasiveness. High density strains are often associated with some detrimental costs for the host: the example of *w*MelPop, as single and triple strain lines in *Ae. albopictus*, shows how strong viral blockers can be pathogenic for the host (41, 42). In this case, the JF.*w*Au line imposes a moderate fitness cost in terms of fecundity and survival rate, when compared to JF, although no difference was observed in the number of eggs laid by individual females when *w*Au is present.

An important factor to be considered with respect to the potential utilization of *w*Au for dengue control is of course the lack of CI induced by this strain. Nevertheless, despite this absence, *w*Au has been shown to be able to rapidly spread and remain at high frequency in *Drosophila simulans* natural populations (31), hypothesised to be a result of providing fitness advantages to its host under some conditions. Further investigation into the ecological circumstances in which fitness benefits could be provided by *w*Au in *Aedes* could offer insights into whether and how this strain could be deployed for population replacement approaches, in light of the very efficient virus blocking it produces.

## Acknowledgments

We thank W. A. Nazni for providing *Ae. albopictus* wild-type eggs. We also thank Giuditta de Lorenzo and Arvind Patel for providing ZIKV antibody.

## Supplementary Information

**Fig S1:**
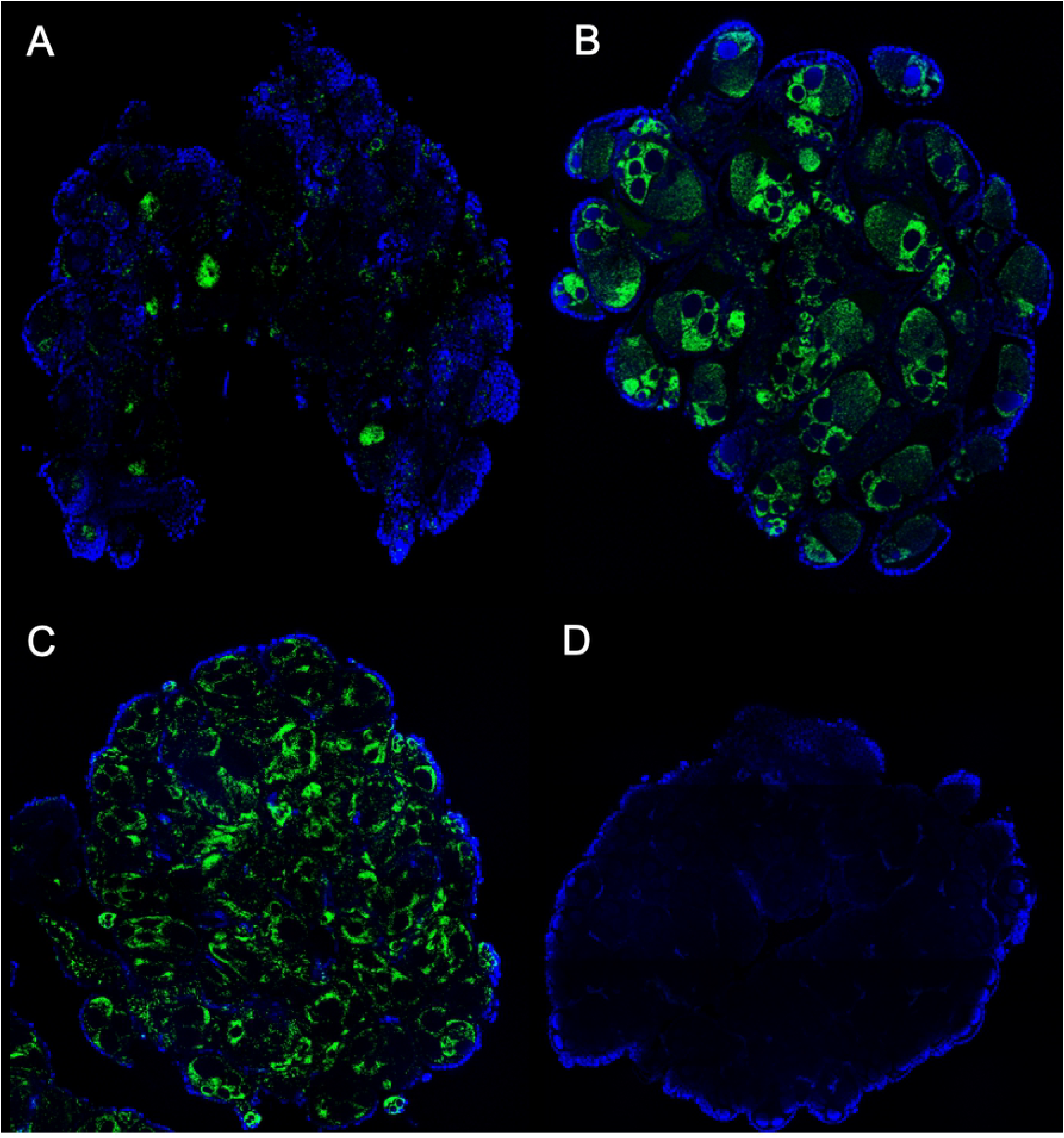
Fluorescent *in situ* hybridization. Visualization of distributions of general *Wolbachia* (green) in ovaries of 5-day old females from the wild-type (A), JF.*w*Au (B), *Ae. aegypti*-*w*Au lines (C) and a no-probe control on wild-type (D). Blue stain is DAPI.

**Table S1:**
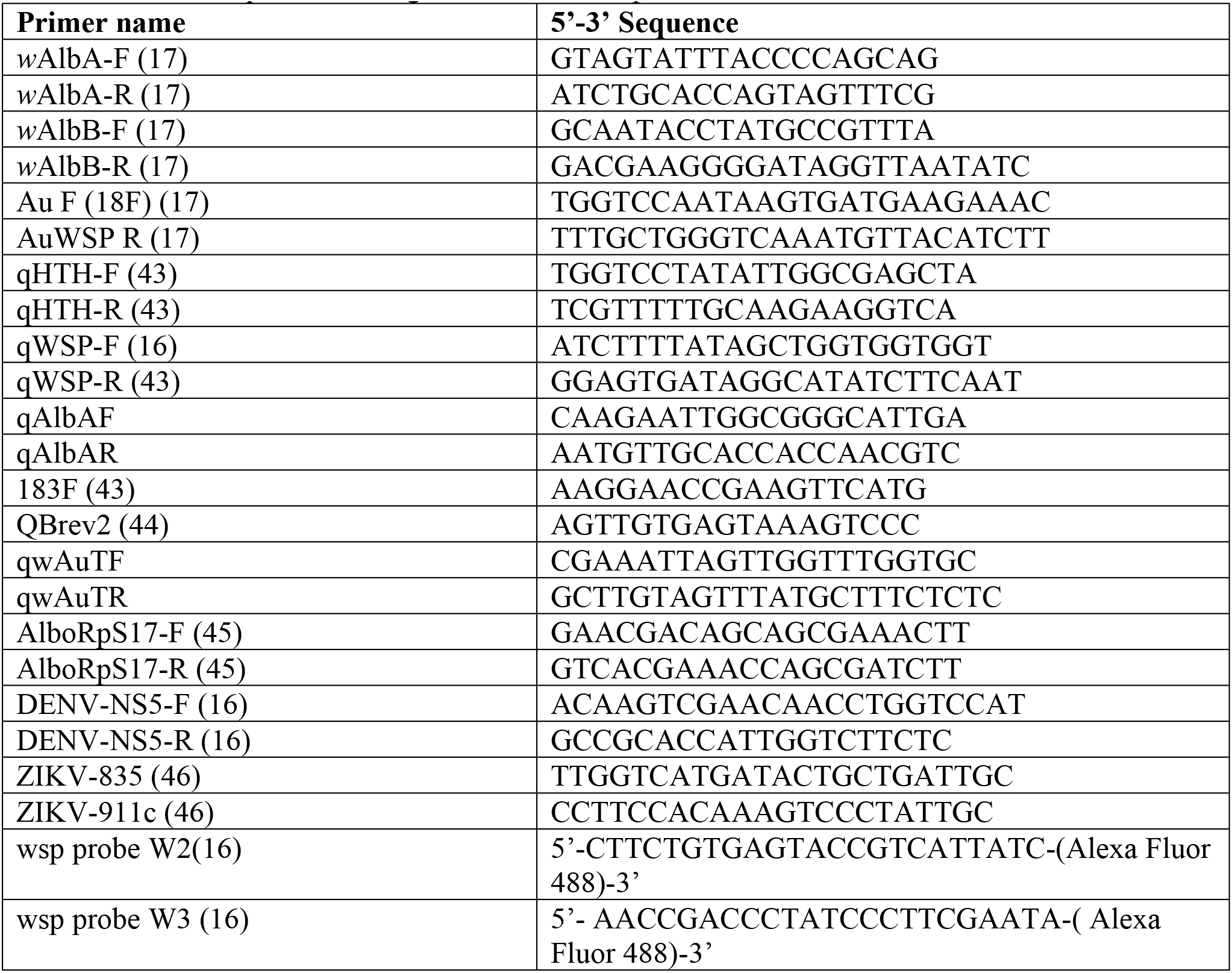
List of sequences of oligonucleotides and probes.

